# Whilst the gymnosperm tree of life may survive the ongoing biodiversity crisis, the angiosperm tree of life may not

**DOI:** 10.1101/2025.03.18.643868

**Authors:** Kowiyou Yessoufou

## Abstract

How would the tree of life be affected by the ongoing biodiversity crisis? We know that the vertebrate tree of life may survive this crisis. However, due to the lack of comprehensive risk assessment for angiosperms, this question could not be fully investigated for vascular plants. Here, I took advantage of the first ever and most recent comprehensive risk predictions for 328 565 angiosperm species to ask whether high-risk vascular plant species represent a disproportionate amount of total evolutionary history. The findings showed significant association between risk status and evolutionary distinctness but only for angiosperm and not for gymnosperm, suggesting that future extinctions may result in a disproportionate pruning of angiosperm tree of life whilst most of the evolutionary history of gymnosperm might persist. To aid bending the curve of loss, we identified most over-threatened clades (e.g., Balsaminaceae) and priority species (e.g., *Hymenophyllum exquisitum*) for conservation.

## Introduction

The world is witnessing an anthropogenically driven biodiversity crisis [1]. In the context of limited resources, conservation efforts must be prioritized [2] with focus on species whose extinction would represent dramatic pruning of the tree of life [3]. The evolutionary distinctness (ED) is now the commonly used metric to identify such species for prioritization purposes or to monitor the tree of life [3,4]. High-ED species may exhibit unique functional traits [5], reveal unique signature of evolution [6], and as such accumulate exceeding evolutionary history [7] representing option values for future ecosystem services [8].

Interestingly, several studies investigated whether ED-informed conservation efforts would effectively protect much of the tree of life (e.g., [9-11]). The common denominator in all these studies is that they all focused on vertebrates, and all revealed that high-ED species are not at higher risk (e.g., reptile, [11]; mammals, [10]; birds, [3]), suggesting that vertebrate tree of life may persist under the current biodiversity crisis. However, we have poor knowledge of whether the same can be said of plants (∼350 000 species). Even in the less diversified gymnosperms (∼1090 species; [12]), we only know the fate of one group, the cycads, which indicates that older cycads (high-ED cycads) are at higher risk, suggesting that much of the cycad tree of life would be disproportionately pruned if all at-risk species slide into extinction [13]. For the most diversified angiosperms, our knowledge of how the ongoing crisis would affect the global tree of life is the poorest due to the lack of a comprehensive threat status for the vast majority of these flowering plants (∼18% of angiosperms have known status, [14], thus preventing the Target 2 of the Global Strategy for Plant Conservation to be achieved [15].

Interestingly, due to improvement in methodological approaches, it is now possible to employ advanced modern techniques such as machine learning to predict the risk status of most species (e.g., [16,17]). Recently, Bachman et al. [17] employed the Bayesian Additive Regression Trees (BART) modelling framework to generate the first comprehensive set of risk status predictions for all flowering plants. Their work finally provided the much needed but long-overdue opportunity to fill out the knowledge gap on the extent of the ongoing crisis on the angiosperm tree of life.

Here, we take advantage of this comprehensive risk status assessment [17] to address the following questions: how would the ongoing biodiversity crisis impact the angiosperm vs. gymnosperm tree of life? Are some clades (e.g., families) over-threatened, that is, do some families contain more threatened species than expected [18]?

## Methods

### Assembling the angiosperm tree of life

The gymnosperm tree of life was retrieved from Forest et al. [12]. To assemble the angiosperm tree of life, we retrieved the latest list of vascular plant species, including 328 565 species, generated in Bachman et al. [17]. This was done as implemented in the R package ‘V.PhyloMaker’ [19]. However, we noted some discrepancies between family names in Bachman et al. [17] and the megatree GBOTB.extended.tre [20,21] available in V.PhyloMaker. These discrepancies were corrected so that family names match that of the megatree. Also, 12 species names were duplicated in the list of Bachman et al. [17], including *Alchemilla sericata, Arctotis decurrens, Bassia aegyptiaca, Celmisia durietzii, Choretrum lateriflorum, Corydalis tenerrima, Helichrysum oligocephalum, Osbeckia capitata, Pterospermum obtusifolium, Rubus newbridgensis, Russelia worthingtonii*, and *Solanum lumholtzianum*. After removing the duplicates, our final dataset includes 328 553 species. The final list of all species was formatted in three columns for species, genus and family names [19], and then, using the R function ‘phylo.maker’, the phylogenetic tree including all species was reconstructed based on the mega-tree ‘GBOTB.extended.tre’ [20,21]. We referred to the resulting angiosperm tree of life (Supplemental information S1).

### Risk status of for gymnosperm and angiosperm species

Gymnosperms risk data were retrieved from Forest et al. [12] (Table S1). In their recent study, Bachman et al. [17] fitted a Bayesian Additive Regression Trees (BART) to range size, human footprint, climate, and evolutionary history of plants data to predict the risk status of all angiosperms and determine the confidence (high vs. low) attached to each prediction. We retrieved their risk predictions (threatened vs. nonthreatened) and analyzed them within a phylogenetic framework (Table S2).

### Identification of over-threatened families

We sorted data by number of threatened species by family and determined total number of species per family (Table S3). We then fitted a negative binomial GLM to the number of threatened species using total number of species in each family as the predictor. We identified over-threatened families as those with positive residuals indicating higher number of threatened species than would be expected from the fit of the model.

### Phylogenetic metrics

We calculated the evolutionary distinctness (ED; [22]; Table S2) for each species using the R function *evol_distinct* in the R library *phyloregion* [23]. Using one-way ANOVA, we tested whether high-ED species (representing much evolutionary history) are at greater risk, a scenario which, if true, might lead to a severe pruning of the tree of life. We did this for both angiosperms and gymnosperms. Next, we determined the EDGE score (Evolutionary Distinctness and Globally Endangered) for each species (EDGE = [ln(1 + ED) + GE*ln(2)], where GE = global endangerment (LC, NT, VU, EN, CR). Since Bachman et al. [17] predicted species risk status as ‘threatened’ vs. ‘nonthreatened’ with high and low confidence, and the calculation of EDGE scores require risk status in the IUCN categories of LC, NT, VU, EN and CR, we adopted the following treatment to Bachman et al.’s predictions: all species predicted with high confidence to be threatened are assigned the IUCN status “CR” whereas those predicted to be threatened with low confidence was assigned “VU”. However, species predicted with high confidence to be nonthreatened were assigned “LC” and those predicted with low confidence to be nonthreatened were assigned “NT”. Similar treatments were done for DD, LR/cd, LR/lc, LR/nt and NE species: they were assigned “CR” when they are predicted with high confidence to be threatened, “VU” when with low confidence, “LC” when predicted as nonthreatened with high confidence and “NT” when predicted as nonthreatened with low confidence (Table S2). Then, we coded GE as follows: LC = 0, NT 249 = 1, VU = 2, EN = 3, CR = 4. We ranked all species based on their EDGE scores to generate a priority list of angiosperms for conservation (Table S4).

### Impacts of biodiversity crisis on the angiosperm tree of Life

We quantified the total phylogenetic diversity PD_total_ on the angiosperm tree of life [24]. Then, we calculated PD_threatened_ corresponding to the amount of PD that would be lost if all threatened species go extinct. Next, we simulated the loss of all threatened species from the tree by pruning randomly the identified number of threatened species 1000 times from the tree. For each random pruning, we calculated the corresponding PD loss termed PDrandom_loss and then compared PD_threatened_ with the average of 1000 PD_random_loss_. All analyses were run in R (see R script in supplemental information S2).

## Results

The total PD on the angiosperm tree of life is PD_total_ = 3 864 572 Myrs. All the threatened species represent PD_threatened_ = 2 250 288 Myrs, corresponding to 58.23% of total PD. This at-risk PD is far greater than expected at random (CI= [1 964 148; 1 977 461]; Figure 1a). Species predicted with high confidence to be threatened represent PD_threatened_confidence_ = 1 929 770 myrs, suggesting that approximately 50% of the total evolutionary history of angiosperms is at higher risk, also exceeding the loss at random (CI = [CI = [1664484; 1678189]; Figure 1b). We calculated the PD already lost due to the extinction of some species (PD_ext_), irrespective of whether they are extinct or extinct in the wild. We found PD_ext_ = 27 662.49 myrs, representing 0.72% of the total PD, and this also exceeds the random loss (CI = [15552.43; 17475.59; Figure 1c).

**Figure 1.**
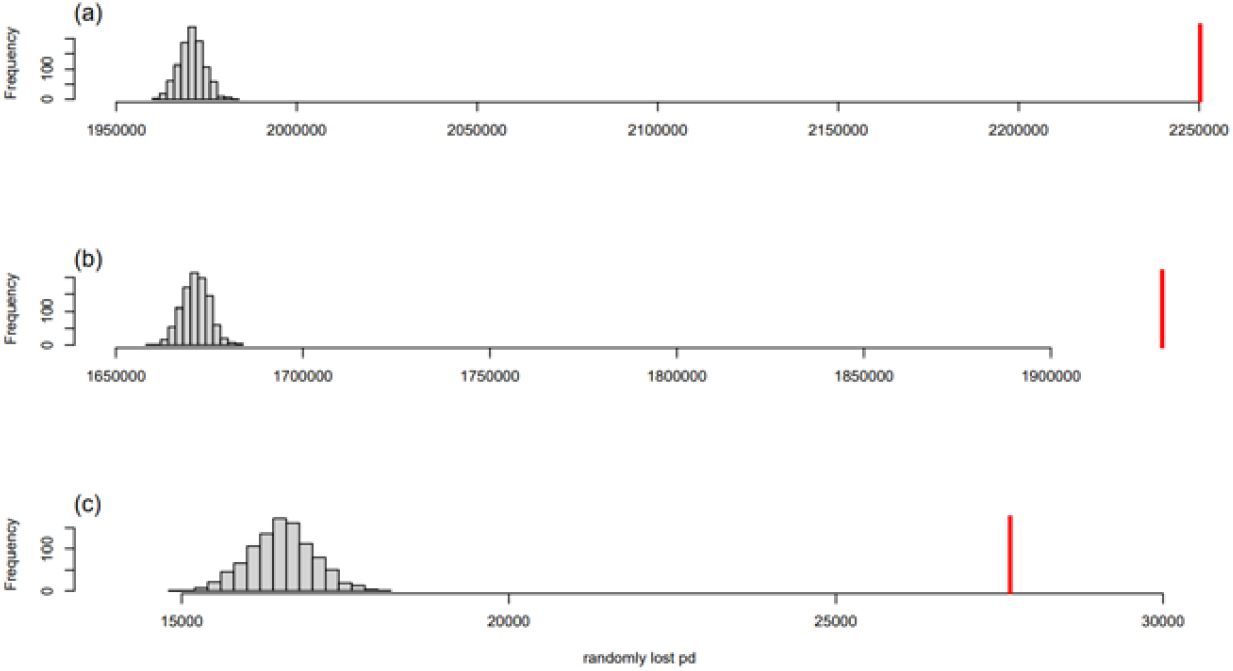
Patterns of PD loss if threatened species go extinct. (a) when all threatened species (actual + predicted to be threatened) are lost, (b) when only species actually threatened + predicted to be threatened with high confidence are lost, and (c) when only extinct species (EW+EX) are pruned.

Furthermore, our negative binomial model indicated that some clades are over-threatened, that is, contain more threatened species than expected (Table S5), and the top 10 of such clades (Figure 2) include Balsaminaceae (model residual = +1.39), Pandanaceae (+1.28), Dipterocarpaceae (+1.24), Berberidaceae (+1.23), Aristolochiaceae (+1.18), Plumbaginaceae (+1.17), Thymelaeaceae (+1.1433), Pentaphylacaceae (+1.1431), Ebenaceae (+1.11), and Actinidiaceae (+1.07).

**Figure 2.**
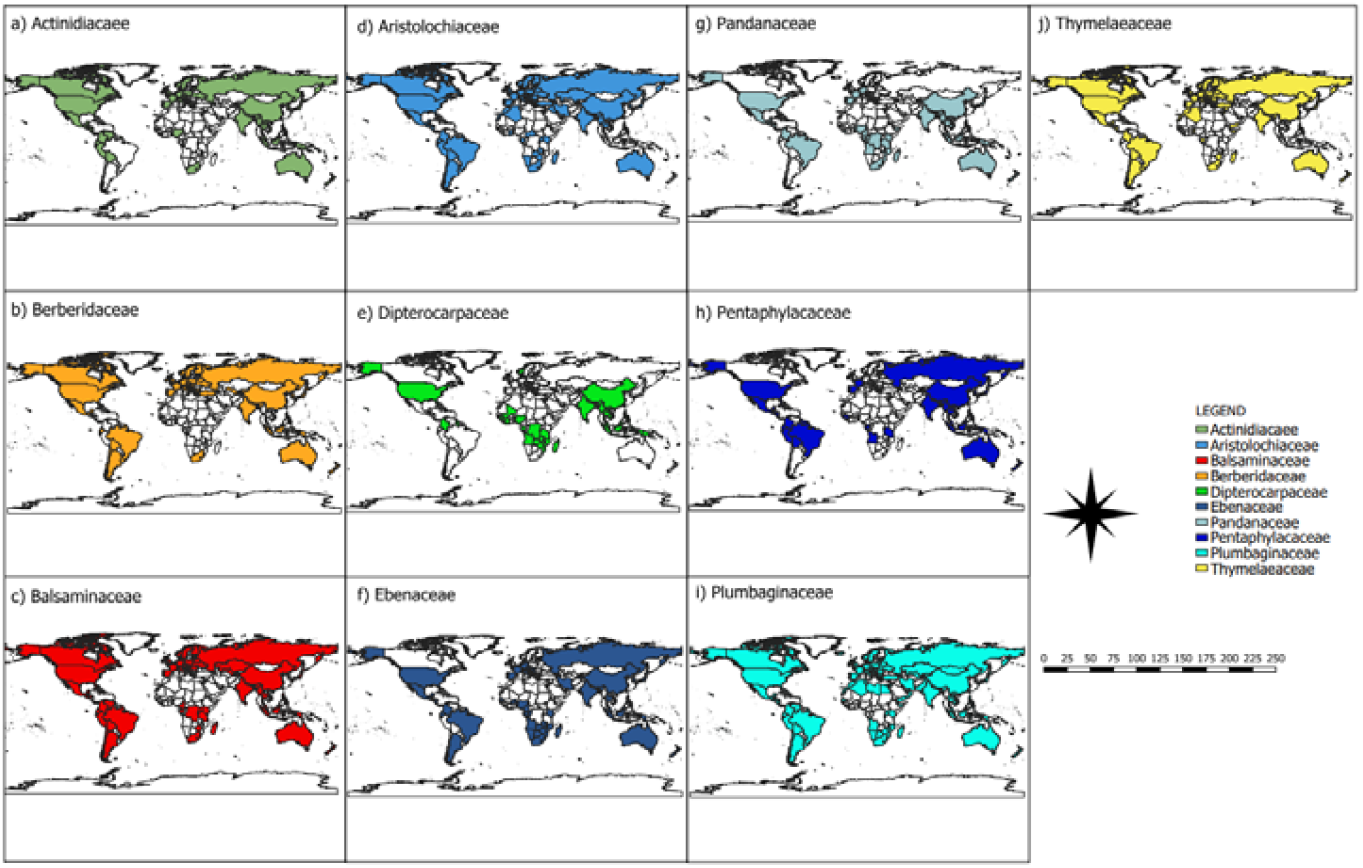
Geography of the top 10 most over-threatened families – families containing more threatened species than expected.

Finally, we used key phylogenetic metrics (ED and EDGE) to aid the prioritization of conservation efforts. We found that evolutionarily unique angiosperm species (high-ED species) are at greater risk whether we considered the status of all species (ANOVA, df = 2, F-value = 555.5, P<0.001; Figure 3a), or distinguished between threatened species vs threatened with high confidence (DF=3, F=371.7, P<0.001; Figure 3b). However, the same cannot be said about gymnosperms (ANOVA, df=2, F-value= 1.722, P=0.179; Figure 3c). We ranked all species based on their EDGE scores (Table S2), and this ranking is indicative of priority list for conservation. The top species in the list is *Hymenophyllum exquisitum* (EDGE = 6.662956756) followed by *Bidayuha crassispatha* (EDGE=5.538390936), and *Englerarum hypnosum* (EDGE= 5.538390936).

**Figure 3.**
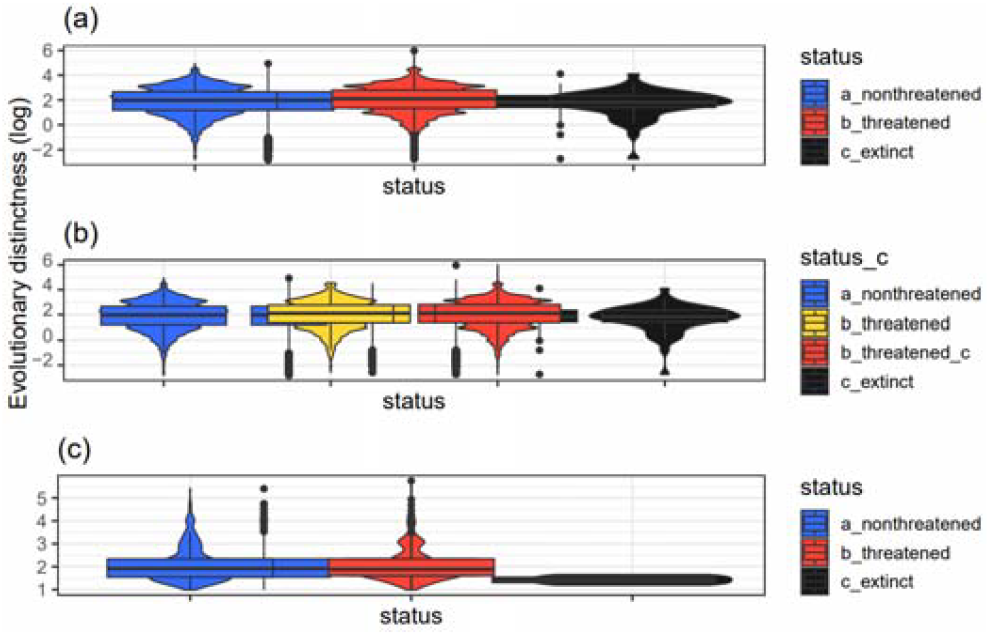
Relationship between risk status and evolutionary distinctness. (a) case of all angiosperm species including all risk status, (b) case of all angiosperm species but with risk status distinguished in threatened vs. threatened with high confidence, and (c) case of all gymnosperms.

## Discussion

An early study estimated the evolutionary history of angiosperm (orders + families) at 35 244 million years [25]. Our finding indicates that the extent of the evolutionary history (PD) of the angiosperm that needs to be preserved is far greater. Unfortunately, as opposed to the 4.1% of this history predicted to be at risk [25], our analysis revealed that it is 58.23% of angiosperm history that may be eroded under current crisis, and this is far greater than expected at random. The pattern of higher loss of PD than expected was found irrespective of whether we distinguished between risk prediction with high confidence or not and even when we analyzed PD already lost due to species already extinct. Only two scenarios can explain such patterns.

First, a large PD loss may be observed when threatened species are evolutionarily distinct. We found indeed that high-ED angiosperm species tend to be threatened. This finding contrasts with what is well established for vertebrates [3,9-11], suggesting that, while vertebrate tree of life may survive the ongoing biodiversity crisis ([26], but see [27]), the angiosperm tree of life may not. However, for gymnosperms, we found that ED does not predict threat status, suggesting that the ongoing extinction crisis may not result in a disproportionate loss of evolutionary history, matching the findings for birds and mammals [3,10] but contrasting with what we found for angiosperm. Our finding also contrasts with that of Vamosi and Wilson [25], who reported that threatened angiosperms are not evolutionarily distinct. This is likely because they assumed in their analysis that every species has a fixed probability of being at risk, while the risk status we used in this study is predicted based on a machine learning approach [17]. In addition, an early regional study found that younger angiosperm clades (low-ED species) tend to be more at risk ([28]; see also [10] for primates), a finding that contradicts ours, suggesting that the global extinction pattern may mask some nuances at regional scale.

Second, a large PD loss may also be observed when threatened species are clustered within some clades ([29,25) but see [30]). Indeed, we found that some families are particularly rich in at-risk species, including Balsaminaceae, Pandanaceae, Dipterocarpaceae, Berberidaceae, Aristolochiaceae, Plumbaginaceae, Thymelaeaceae, Pentaphylacaceae, Ebenaceae, and Actinidiaceae. The consequence of such risk patterns is that the ongoing biodiversity crisis may result in the loss of entire clades (see also [25]). Indeed, according to Vamosi and Wilson [25], one-third of at-risk species may be lost in the next 100 years. However, it is important to highlight that the loss of large PD is contingent upon tree shape and a pruning of high-ED clades [30]. We suggest that the large PD loss may rather be the result of the pruning of greater number of branches due to a non-random extinction that might preferentially target some clades [27].

Finally, to shed light on which keys to press to bend the curve of biodiversity loss and accelerate recovery [4,31-33], I ranked species based on their EDGE scores. The top species in the list is *Hymenophyllum exquisitum* (EDGE = 6.662956756) which is native to Taiwan and found in the subtropical biome at high elevations (1200-2700m asl) growing most of the time as epiphytes. This is followed by *Bidayuha crassispatha* (EDGE=5.538390936), native to Borneo and growing in the wet tropical biome. The third most deserving species for conservation is the perennial or subshrub *Englerarum hypnosum* (EDGE= 5.538390936) distributed from China to Indo-China in subtropical biome.

For far too long, we had no comprehensive insight into the risk status of all angiosperms (∼350 000 species) as opposed to vertebrates. This prevents conservation scientists from investigating the extent of the consequences of the ongoing biodiversity crisis on the angiosperm tree of life. Bachman et al. [17] recently provided, using machine learning, this missing information on angiosperm. I took advantage of their work to show that the evolutionary history to be lost exceeds the expectations at random, and that the angiosperm tree of life may not survive the crisis unlike the gymnosperm. This finding implies that there is a need to renew global efforts and commitments to bend the curve of biodiversity loss. Future works must collect and analyze additional data on spatial, geographic, ecological, and life-history of angiosperms to identify clades and regions of exceptionally high conservation priority [11].

## Ethics

This work did not require ethical approval from a human subject or animal welfare committee.

## Data accessibility

The data used were retrieved from Bachman et al. (2024). However, we applied some treatments and the resulting data and R scripts that support the findings of this study are available at ref. [34] (Figshare, Dataset, https://figshare.com/s/32a4891bdf9eb22e8fd6).

## Declaration of AI use

I did not use AI

## Authors’ contributions

K.Y.: conceptualization, data curation, formal analysis, investigation, methodology, project administration, software, validation, visualization, writing—original draft, writing—review and editing. I gave final approval for publication and agreed to be held accountable for the work performed therein.

## Acknowledgements

I thank the National Research Foundation— South Africa for financial support.

## Supplemental Information

**Table S1**. Gymnosperms data on ED and EDGE scores, retrieved from Forest et al. [12].

**Table S2**. Risk status retrieved from Bachman et al. [17]. Observed threat = observed risk status; iucn_category_treated = Species IUCN status determined inferred from Bachman et al. [17] predictions as explained in the method; confidence: confident = risk status of corresponding species is predicted with high confidence; low confidence = status predicted with low confidence (extracted from Bachman et al. [17]); status = risk status as predicted by Bachman et al. [17]; status_c = risk status as predicted by Bachman et al. [17] but with highlight of risk predicted with high confidence (c for figh confidence); ED = evolutionary distinctness

**Table S3**. Summary of species risk status per family. tot_species = total number of species per family, predicted_threat = risk status as predicted by Bachman et al. [17]; threatened = number of threatened species per family. This is the dataframe used for our negative binomial model (see materials and method section)

**Table S4**. Angiosperm species ranked based on their EDGE scores.

**Table S5**. Family ranking based on residuals following fitting negative binomial model. resid_nb = values of residuals ranked from the highest (corresponding to the most over-threatened family) to the lowest (corresponding to the under-threatened family).

**S1**. Angiosperm tree of life

**S2**. R scripts used to analyze all the data

