## Supplementary material for "Whilst the gymnosperm tree of life may survive the ongoing biodiversity crisis, the angiosperm tree of life may not": R script_global_Extinction risk.R

```
#analysis of extinction risk of global angiosperm
#by Prof. Kowiyou Yessoufou

rm(list=ls(all=TRUE))

# Taxonomic analysis

#Question 1: What are hot nodes of threatened species? 
  
## A. Dataset
data_ext_global_family <- read.table("C:\\Users\\kowiyouy\\OneDrive - University of Johannesburg\\Desktop\\Kowiyou\\my papers\\DATABASES\\ALL angiosperms risk predictions\\data\\paper 1\\data\\Angiosperm_sorted by family.txt",header = TRUE)
attach(data_ext_global_family)
names(data_ext_global_family)

## Sort data in ascending order to facilitate plotting credible intervals later in the code.

data_ext_global_family<-data_ext_global_family[order(data_ext_global_family$tot_species),]

## Checking the nature of each variable
str(data_ext_global_family)

attach(data_ext_global_family)
head(data_ext_global_family)

library(MASS)
## install.packages("calibrate")
library(calibrate) ## add label to data point

## fitting poisson and negative binomial

## We fit a poisson model first
poim<-glm(threatened~tot_species, poisson)
summary(poim)

## there is a strong overdispersion, deviance/df= 658869/414 = 1591.471 >>>>1. We will use negative binomial error structure instead to account for overdispersion

poi_nb<-glm.nb(threatened~tot_species)
summary(poi_nb)

# 
#            Estimate   Std. Error  z value  Pr(>|z|)    
#tot_species 1.222e-03  3.223e-05   37.91   <2e-16 ***
#  ---

## This suggest a significant positive link between family size and number of threatened plant species:
#. beta=1.222e-03 +/- 3.223e-05 (Z=37.91, p<0.00001, AIC: 4322.3). 
#The mean number of threatened species is 1.001=exp(1.222e-03) per family.

## The residuals are:
predict_nb<-predict(poi_nb,  type="response")
resid_nb<-residuals(poi_nb)
resid_nb_family<-data.frame(family, resid_nb)
write.csv(resid_nb_family, "global_Residual_NegBin.csv")


#phylogenetic analysis
# load the package
library(ape)
#library("V.PhyloMaker")
#library(utf8)


# form the species list for all the species
species_list <- read.table("C:\\Users\\kowiyouy\\OneDrive - University of Johannesburg\\Desktop\\Kowiyou\\my papers\\DATABASES\\ALL angiosperms risk predictions\\data\\paper 1\\data\\species list.txt",header=TRUE)
# generate the phylogeny presented in
#species_list$species
#utf8_normalize(species_list$species)
#as_utf8(species_list$species, normalize = TRUE)
#as_utf8(species_list$species)
#utf8_valid(species_list$species)

tree.life <- phylo.maker(sp.list = species_list,
                           tree = GBOTB.extended, nodes = nodes.info.1, scenarios="S3")
tree.life <- tree.life$scenario.3

write.nexus(tree.life, file = "tree_life_global")


#plot(tree.life,cex=0.001)

# call the tree
library(phyloregion)
tree.life <- read.nexus("C:\\Users\\kowiyouy\\OneDrive - University of Johannesburg\\Desktop\\Kowiyou\\my papers\\DATABASES\\ALL angiosperms risk predictions\\data\\paper 1\\data\\tree_life_global.txt")

# calculate ED (evolutionary distinctiveness)
ed_global <- evol_distinct(tree.life, type = "fair.proportion")

#save ED values
write.table(ed_global, file = "ed_global")

# Analysis of extinction risk
#question 1: Are threatened angiosperm species evolutionary more distinct than not?

data_ext_global <- read.table("C:\\Users\\kowiyouy\\OneDrive - University of Johannesburg\\Desktop\\Kowiyou\\my papers\\DATABASES\\ALL angiosperms risk predictions\\data\\paper 1\\data\\Angiosperm_extinction_risk_predictions_v1.txt",header=TRUE)
attach(data_ext_global)
names(data_ext_global)

#fitting models and graphically represent the relationships between ED and IUCN status

model1 <- aov(log(ED)~status,data=data_ext_global)
summary(model1)
#                 Df    Sum Sq  MeanSq  Fvalue Pr(>F)    
#status           2     1309    654.6   555.5  <2e-16 ***# high-ED angiosperms are at greater risk
#Residuals       328551 387150  1.2  

op<-par(mfrow=c(2,2), pty="s", las=1)

boxplot(log(ED)~status,data=data_ext_global,main="",notch=TRUE,xlab="",ylab="Evolutionary distinctiveness (log)",col=c("blue","red","black"),names=c("nonthreatened","threatened","extinct"))
mtext("(a)", side=3, adj=0)


#same analysis but distinguishing between threat prediction with high confidence vs prediction with low confidence
model1_c <- aov(log(ED)~status_c,data=data_ext_global)
summary(model1_c)
#             Df     Sum Sq  Mean Sq  F value Pr(>F)    
# status_c    3      1314    438.0    371.7   <2e-16 ***
# Residuals   328550 387145  1.2 

boxplot(log(ED)~status_c,data=data_ext_global,main="",notch=TRUE,xlab="",ylab="",col=c("blue","orange","red","black"),names=c("nonthreatened","threatened","threatened_c","extinct"))
mtext("(b)", side=3, adj=0)


#question 2: Are threatened gymnosperm species evolutionary more distinct than not?

data_ext_global_gymn <- read.table("C:\\Users\\kowiyouy\\OneDrive - University of Johannesburg\\Desktop\\Kowiyou\\my papers\\DATABASES\\ALL angiosperms risk predictions\\data\\paper 1\\data\\gymnosperms\\data_gymnopserms on the edge.txt",header=TRUE)
attach(data_ext_global_gymn)
names(data_ext_global_gymn)

#same analysis assuming all DD and NE are threatened (see Forest et al. 2018 Gymnosperms in the edge. Scientific Report 8:6053)
model3 <- aov(log(ED)~status,data=data_ext_global_gymn)
summary(model3)
#                  Df  Sum Sq  Mean Sq  F value Pr(>F)
#threat_binary     2    1.8    0.9056   1.722   0.179 # high-ED gymnosperms are not at greater risk 
#Residuals        1087  571.7  0.5260 

boxplot(log(ED)~status,data=data_ext_global_gymn,main="",notch=TRUE,xlab="threat categories",ylab="Evolutionary distintiveness",col=c("blue","red","black"),names=c("nonthreatened","threatened","extinct"))
mtext("(c)", side=3, adj=0)

#All 3 figures
#par(mfrow=c(2,2))
library(tidyverse)
library(RColorBrewer)

Fig2a<-ggplot(data_ext_global,aes(x=status,
                           y=log(ED),
                           fill=status))+
  geom_violin()+
  geom_boxplot(width=2)+
  geom_boxplot(show.legend = FALSE,
               outlier.shape = NA)+
  labs(title = "(a)", x='status', y="") +
  theme_bw(14)+
  theme(
    axis.text.x=element_blank())+
  scale_fill_manual(values=
                       c("#336BFF",
                         "#FF4233",
                         "#161719"))
                   
###
Fig2b<-ggplot(data_ext_global,aes(x=status_c,
                                  y=log(ED),
                                  fill=status_c))+
  geom_violin()+
  geom_boxplot(width=2)+
  geom_boxplot(show.legend = FALSE,
               outlier.shape = NA)+
  labs(title = "(b)", x='status', y="Evolutionary distinctness (log)") +
  theme_bw(14)+
  theme(
    axis.text.y = element_text(vjust = 0.1),
    axis.title.y = element_text(hjust = 0.15),
    axis.text.x=element_blank())+
  scale_fill_manual(values=
                      c("#336BFF",
                        "#FFD733",
                        "#FF4233",
                        "#161719"))

#
###
Fig2c<-ggplot(data_ext_global_gymn,aes(x=status,
                                  y=log(ED),
                                  fill=status))+
  geom_violin()+
  geom_boxplot(width=2)+
  geom_boxplot(show.legend = FALSE,
               outlier.shape = NA)+
  labs(title = "(c)", x='status', y="") +
  theme_bw(14)+
  theme(
    axis.text.x=element_blank())+
  scale_fill_manual(values=
                      c("#336BFF",
                        "#FF4233",
                        "#161719"))

## Fig 2 ***************************
gridExtra::grid.arrange(Fig2a, Fig2b, Fig2c, nrow=3)


#Angiosperms on the EDGE: Creating a list of ranked angiosperm based on EDGE score
library(phyloregion)
edge_angiosperm <- EDGE(data_ext_global,tree.life, Redlist="iucn_category_treated",species="species")
write.table(edge_angiosperm, file = "edge_angiosperm")

###Question 3: Are we going to lose more PD than at random if all all threatened species go extinct?


# Calculate total PD on the phylogenetic tree
total_pd <- sum(tree.life$edge.length) # total_pd = 3864572

#form the list of all threatened species

#firstly, create the list of nonthreatened species
list_nonthreatened <- data_ext_global$species[!data_ext_global$status=="b_threatened"]

#secondly, drop all nonthreatened species from the tree to have a tree of only threatened species
extinct.tree <- drop.tip(tree.life,as.vector(list_nonthreatened))

# Calculate threatened PD i.e. total PD on the tree containing only threatened species
pd_threatened <- sum(extinct.tree$edge.length) #2 250 288

#Calculate the proportion of threatened pd

100*pd_threatened/total_pd # 0.6841% 

#Investigating how a random loss of a number of species equivalent to the number of threatened species would impact the tree
# This requires the following simulation of species loss (same number of species as real number of threatened species are pruned randomly 1000 times from the tree)

# calculate number of threatened species on the phylogeny
number_threatened_species <- length(extinct.tree$tip.label)

#number_threatened_species= 176307

# simulation of loss of 176307 species pruned randomly 1,000 times from the tree, and calculation of pd remaining on the tree each time after a random loss of  175600 species
pd <- NULL
for(x in 1:1000){
  rand.tree <- drop.tip(tree.life,sample(tree.life$tip.label,number_threatened_species))
  pd[x] <- sum(rand.tree$edge.length)
}

# Calculation of the pd corresponding to each set of 175600 species randomly lost from the tree
pd_random_loss <- total_pd-pd # there are 1,000 values of pd_random_loss
mean(pd_random_loss) #1970980
#plot the results in a form of histogram
par(mfrow=c(3,1))

hist(total_pd-pd,main="",xlim = c(1950000,2240000),xlab="") # histogram of random loss
mtext("(a)", side=3, adj=0)

abline(v=pd_threatened,col="red",lwd=3) # v= 2 250 288; vertical line in red showing actual loss
#abline(v=mean(total_pd-pd),col="blue",lwd=3)# mean random pd loss = 139083.2; vertical line in blue showing the mean of random loss
CI <- quantile(total_pd-pd,prob=c(0.025,0.975)) # confidence interval
#CI = [1964148; 1977461] 

######
######################################

# Run the same analysis for species predicted with confidence to be threatened
#form the list of all threatened species predicted with high confidence

#firstly, create the list of species excluding those that predicted as threatened with confidence
list_nonthreatened_c <- data_ext_global$species[!data_ext_global$status_c=="b_threatened_c"]

#secondly, drop all nonthreatened species from the tree to have a tree of only species predicted with confidence to be threatened  
extinct.tree_c <- drop.tip(tree.life,as.vector(list_nonthreatened_c))

# Calculate threatened PD i.e. total PD on the tree containing only species predicted with confidence to be threatened 
pd_threatened_c <- sum(extinct.tree_c$edge.length) # 1929770

#Calculate the proportion of threatened pd with confidence

100*pd_threatened_c/total_pd # = 49.93488 ~50%

#Investigating how a random loss of a number of species equivalent to the number of threatened species would impact the tree
# This requires the following simulation of species loss (same number of species as real number of threatened species are pruned randomly 1000 times from the tree)

# calculate number of species predicted with confidence to be threatened  on the phylogeny
number_threatened_species_c <- length(extinct.tree_c$tip.label)

#number_threatened_species_c= 149872

# simulation of loss of 149066 species pruned randomly 1,000 times from the tree, and calculation of pd remaining on the tree each time after a random loss of 12647 species
pd_c <- NULL
for(x in 1:1000){
  rand.tree <- drop.tip(tree.life,sample(tree.life$tip.label,number_threatened_species_c))
  pd_c[x] <- sum(rand.tree$edge.length)
}

# Calculation of the pd corresponding to each set of xxx species randomly lost from the tree
pd_random_loss_c <- total_pd-pd_c # there are 1,000 values of pd_random_loss
mean(pd_random_loss_c) # 1 671 377

#plot the results in a form of histogram

hist(total_pd-pd_c,main="",xlim = c(1650092,1930000),xlab="") # histogram of random loss
mtext("(b)", side=3, adj=0)

abline(v=pd_threatened_c,col="red",lwd=3) # v= 179194.2; vertical line in red showing actual loss
#abline(v=mean(total_pd-pd_c),col="blue",lwd=3)# mean random pd loss = 139083.2; vertical line in blue showing the mean of random loss
CI <- quantile(total_pd-pd_c,prob=c(0.025,0.975)) # confidence interval
#CI = [1664484; 1678189 ] 


########################
# Run the same analysis for species that are already extinct (EX+EW)
# form the list of all extinct species 

#firstly, create the list of extant species
list_extant <- data_ext_global$species[!data_ext_global$status_c=="c_extinct"]

#secondly, drop all nonthreatened species from the tree to have a tree of only species predicted with confidence to be threatened  
extinct.tree_extinct <- drop.tip(tree.life,as.vector(list_extant))

# Calculate threatened PD i.e. total PD on the tree containing only species predicted with confidence to be threatened 
pd_extinct <- sum(extinct.tree_extinct$edge.length) # 27662.49

#Calculate the proportion of threatened pd with confidence

100*pd_extinct/total_pd # = 0.72%

#Investigating how a random loss of a number of species equivalent to the number of threatened species would impact the tree
# This requires the following simulation of species loss (same number of species as real number of threatened species are pruned randomly 1000 times from the tree)

# calculate number of species predicted with confidence to be threatened  on the phylogeny
number_extinct <- length(extinct.tree_extinct$tip.label) #1494

#number_threatened_species_c= 1494

# simulation of loss of 49 species pruned randomly 1,000 times from the tree, and calculation of pd remaining on the tree each time after a random loss of 12647 species
pd_ext <- NULL
for(x in 1:1000){
  rand.tree <- drop.tip(tree.life,sample(tree.life$tip.label,number_extinct))
  pd_ext[x] <- sum(rand.tree$edge.length)
}

# Calculation of the pd corresponding to each set of 1494 species randomly lost from the tree
pd_random_loss_ext <- total_pd-pd_ext # there are 1,000 values of pd_random_loss
mean(pd_random_loss_ext) #16528.27

#plot the results in a form of histogram

hist(total_pd-pd_ext,main="",xlim = c(14000,30000),xlab="randomly lost pd") # histogram of random loss
mtext("(c)", side=3, adj=0)

abline(v=pd_extinct,col="red",lwd=3) # v= 179194.2; vertical line in red showing actual loss
#abline(v=mean(total_pd-pd_c),col="blue",lwd=3)# mean random pd loss = 139083.2; vertical line in blue showing the mean of random loss
CI <- quantile(total_pd-pd_ext,prob=c(0.025,0.975)) # confidence interval
#CI = [15552.43; 17475.59] 

#END
```
